# Heterogeneous habenular neuronal ensembles during selection of defensive behaviors

**DOI:** 10.1101/2019.12.22.886457

**Authors:** Salvatore Lecca, Vijay MK Namboodiri, Leonardo Restivo, Nicolas Gervasi, Giuliano Pillolla, Garret D. Stuber, Manuel Mameli

**Affiliations:** The Department of Fundamental Neuroscience, The University of Lausanne 1005 Lausanne, Switzerland; Center for the Neurobiology of Addiction, Pain, and Emotion, Department of Anesthesiology and Pain Medicine, Department of Pharmacology, University of Washington, Seattle, WA, USA; College de France, Inserm, 75005 Paris, France; The University of Cagliari, Cagliari, Italy; Inserm, UMR-S 839, 75005 Paris, France

**Keywords:** Lateral habenula, defensive behaviors, single cell calcium imaging in vivo, aversion

## Abstract

Optimal selection of threat-driven defensive behaviors is paramount to an animal’s survival. The lateral habenula (LHb) is a key neuronal hub coordinating behavioral responses to aversive stimuli. Yet, how individual LHb neurons represent defensive behaviors in response to threats remains unknown. Here we show that, in mice, a visual threat promotes distinct defensive behaviors, namely runaway (escape) and action-locking (immobile-like). Fiber photometry of bulk LHb neuronal activity in behaving animals revealed an increase and decrease of calcium signal time-locked with runaway and action-locking, respectively. Imaging single-cell calcium dynamics across distinct threat-driven behaviors identified independently active LHb neuronal clusters. These clusters participate during specific time epochs of defensive behaviors. Decoding analysis of this neuronal activity unveiled that some LHb clusters either predict the upcoming selection of the defensive action or represent the selected action. Thus, heterogeneous neuronal clusters in LHb predict or reflect the selection of distinct threat-driven defensive behaviors.

## Introduction

When facing an external threat, animals select from a repertoire of innate behavioral responses ranging from escape (runaway) to immobile-like (action-locking) strategies (Evans et al., 2019). These behaviors ultimately increase individual survival, rely on the external environment, and can be adopted by the same animal (De Franceschi et al., 2016; Eilam, 2005). The detection of a threat and the optimal selection of such threat-driven actions (i.e. runaway or action-locking) require the coordination of complex brain networks. The recent analysis of threat-driven escape behaviors unraveled the essential contribution of neuronal circuits including the amygdala, the superior colliculus, the periaqueductal grey, the hypothalamus or the midbrain. All of these are pivotal neuronal nodes for aversive processing (Evans et al., 2018; Headley et al., 2019; Silva et al., 2016; Tovote et al., 2016; Zhou et al., 2019). Neurons located in the epithalamic lateral habenula (LHb) signal the negative valence of a stimulus contributing to aversive behaviors (Matsumoto and Hikosaka, 2007). Accordingly, habenular neurons in fish, rodents and non-human primates, as opposed to midbrain dopamine neurons, respond mainly with an excitation to a variety of aversive stimuli, and reduce their activity after reward presentation (Andalman et al., 2019; Lecca et al., 2017; Matsumoto and Hikosaka, 2007; Wang et al., 2017). Specifically, aversion-driven LHb neuronal excitation requires hypothalamic glutamate release to shape behavioral responses upon unexpected and predicted aversive events (Lazaridis et al., 2019; Lecca et al., 2017; Trusel et al., 2019). Indeed, reducing the efficacy of hypothalamus-to-LHb projections impairs behavioral escape driven by foot shocks, shock-predicting cues and predator-like looming stimulus (Lecca et al., 2017; Trusel et al., 2019). The latter evidence indicates a relevant contribution of LHb in encoding environmental threats. Yet, whether specific neuronal representations in the LHb participate in the selection of threat-driven defensive behaviors (runaway or action-locking), remains unknown.

To examine this question, we performed deep-brain Ca^2+^-imaging of large LHb neuronal populations using a head-mounted miniaturized microscope in mice engaging visual threat-driven defensive responses (Resendez et al., 2016). We combined such large-scale recordings with unsupervised classification of response patterns. This led to the identification of functionally distinct LHb neuronal subpopulations during threat-driven runaway and action-locking. Analysis of responses indicates that multiple neuronal clusters emerge during behavioral strategies holding independent information (i.e. prediction vs action) related to the temporal expression of the behaviors. Altogether, these data support the participation of LHb neuronal populations in the selection of defensive behaviors when facing an external threat.

## Results

### Opposing behavioral strategies in response to a visual threat

Ethological studies posit a relationship between the animal-nest distance and the strategy adopted to react to a threat. The closer to a nest, the more likely it is for animals to rapidly runaway to hide. Action-locking responses, instead, occur with higher frequency when the animal is located far from the shelter (Yilmaz and Meister, 2013).

Here we investigated these independent threat-driven behavioral strategies in mice using an innately aversive overhead expanding spot (Looming) (Yilmaz and Meister, 2013), while mice explore an experimental arena provided with a nest. We randomly triggered the looming stimulus when the mouse explored different zones of the arena with variable distance with respect to the nest (Figure 1A and B). Mice predominantly adopted threat-driven high-speed runaway responses (Figure 1A-D). In a smaller fraction of trials, however, the same animals engaged in a looming-driven action-locking, a behavior outlined by significant speed reduction (Figure 1A-D). Such opposite threat-driven behavioral strategies related to the distance from the nest (Figure 1D).

**Figure 1.**
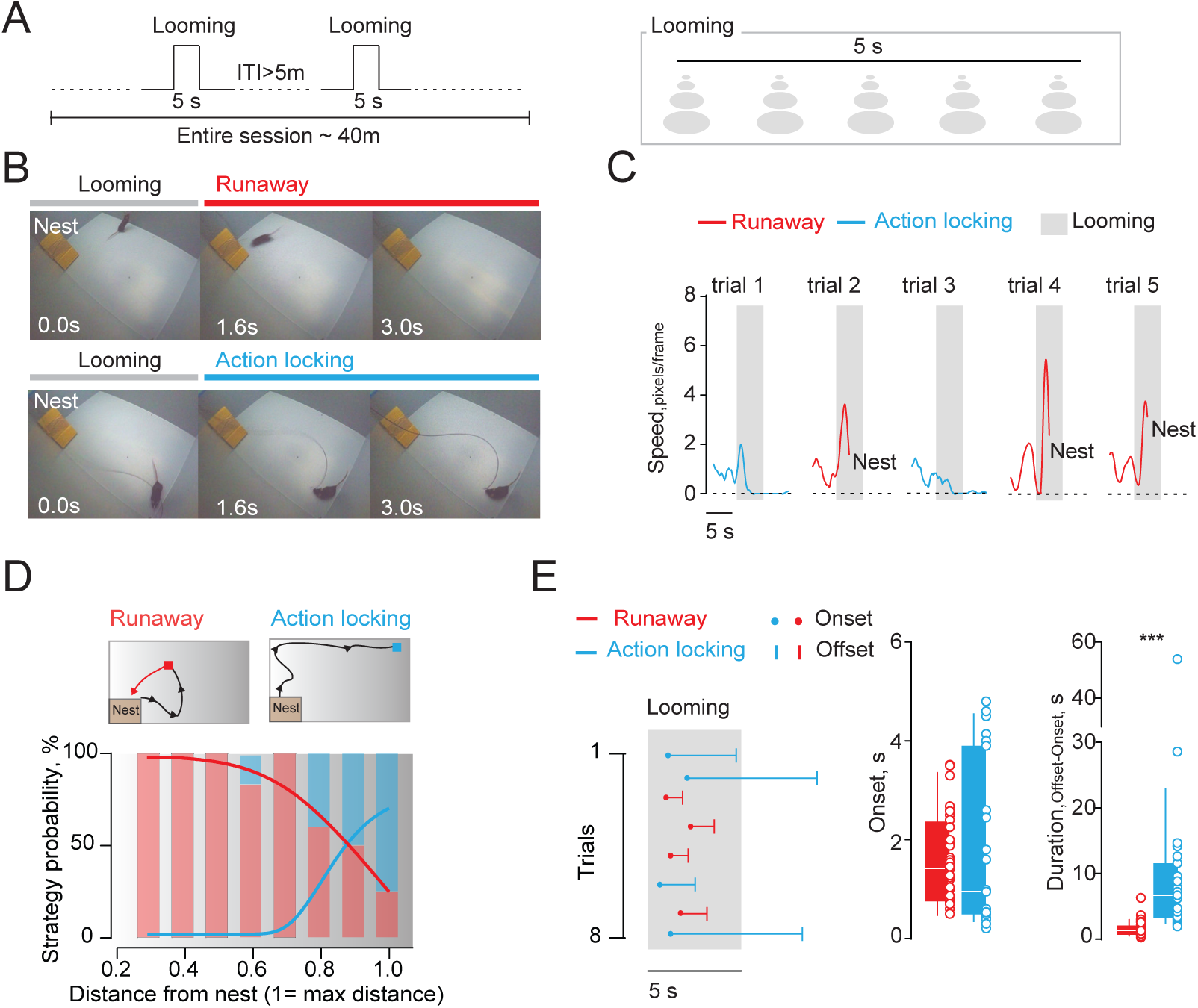
Threat exposure promotes divergent defensive strategies. (A) Schematic of the looming protocol. (B) Extracted video frames depicting a mouse during looming-driven runaway (top) and action-locking (bottom). (C) Representative single mouse runaway and action-locking trials to multiple looming stimuli. (D) Top: representative track of a single mouse during a runaway and an action-locking trials. Bottom: strategy probability in function of the mouse-nest distance (n_runaway trials_= 56; n_action-locking trials_= 23; n_mice_= 11; *Mouse-nest distance (Max distance = 1)*: R trials vs AL trials; *0.3:* 4 vs 0; *0.4:* 7 vs 0; *0.5*: 8 vs 0; *0.6*: 10 vs 2; *0.7:* 11 vs 0; *0.8*: 9 vs 6; *0.9*: 3 vs 3; *1.0*: 4 vs 12; X^2^_7_ = 31.68; ***p<0.0001, Chi Square test). The lines fitting a sigmoidal distribution reports the correlation between the mouse-nest distance and the selected strategy (Runaway: r=-0.883, R^2^=0.78, **p=0.003; Action-locking: r=0.884, R^2^=0.78**p=0.003, Pearson correlation coefficient) (E) Left: Single mouse runaway (R, in red) and action-locking (AL, in blue) timeframe reported for each trial (dot: onset response, line: offset response). Right: pooled data (n_runaway trials_= 56; n_action-locking trials_= 23) for onset (R vs AL; 1.631±0.14 vs 1.797±0.34 s; t_77_= 0.53; p= 0.59, unpaired t-test) and duration (R vs AL; 1.52 ± 0.14 vs 9.58 ± 2.37 s; t_77_=5.29; ***p<0.0001, unpaired t-test) of runaway and action-locking. Data are presented with boxplots (median and 10-90 quartile) or mean ± S.E.M.

Multiple looming presentations (maximum of 12) revealed comparable average onset time between runaway and action-locking responses, yet different offset timing, with action-locking events lasting up to tens of seconds (Figure 1E). Altogether, mice can display divergent defensive behaviors to the same visual threat stimulus in a context-dependent fashion.

### Threat encoding in the lateral habenula

We next employed fiber photometry to measure fluorescent calcium transients (Ca^2+^; (Cui et al., 2014)) and examined the population dynamics of LHb neurons in freely behaving mice (Figure 2A). We injected rAAV2.5-hSyn1-GCaMP6f into the LHb and implanted an optical fiber above the injection site (Figure 2A and Figure 2– figure supplement 1A). The onset of threat-driven runaway occurred along with a robust increase in Ca^2+^ fluorescence from LHb neurons (Figure 2A, B and movie 1). In contrast, looming-driven action-locking developed together with a significant reduction in LHb fluorescence (Figure 2A, B and movie 2). Notably, a significant shift in fluorescence emerged time-locked with the visual looming stimulus and prior the behavior (Figure 2–figure supplement 1B, C). The magnitude of this fluorescence rise was comparable between runaway and action-locking trials (Figure 2–figure supplement 1B, C). The observation that no fluorescence transients occurred in animals injected only with a rAAV2.5-hSyn1-GFP, supports the specificity of signal detection (Figure 2–figure supplement 2A, B). Both runaway and action-locking expressed along with an abrupt change in speed at the behavioral onset (Figure 1C). However, speed changes outside the looming presentation did not coincide with fluorescence transients, supporting that spontaneous locomotion does not engage LHb activity (Figure 2–figure supplement 3A, B; (Lecca et al., 2017)). Altogether, these data suggest that a threat recruits differential LHb neuronal responses throughout the expression of diverse behavioral strategies (i.e. from stimulus detection to action completion).

**Figure 2.**
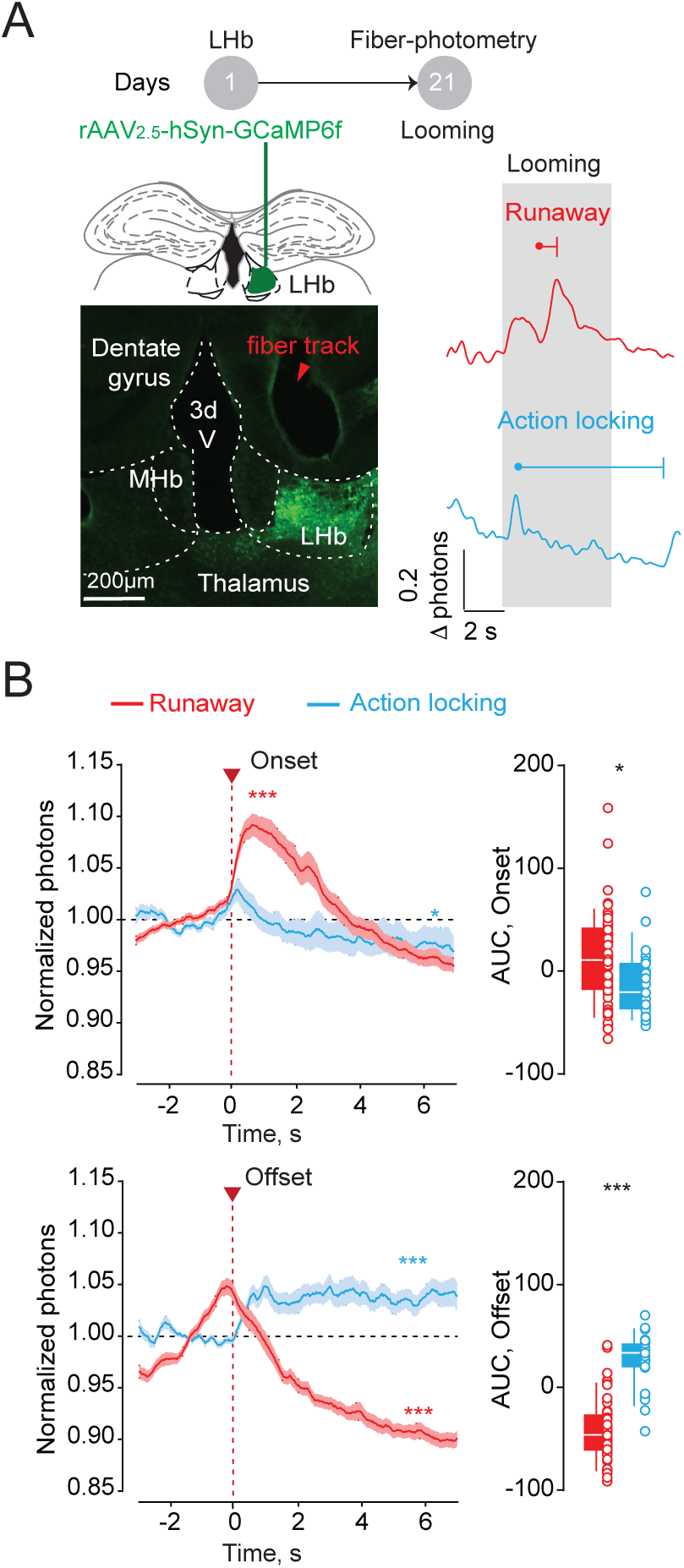
Opposite habenular neuronal dynamics during divergent defensive strategies. (A) Top: schematic of the experiment. Bottom left: representative brain coronal section showing GCamp6f transduction and the fiber implantation track in the LHb. Bottom right: representative Ca^2+^ traces during runaway (R, red, top) and action locking (AL, blue, bottom) trials (Looming, gray bar). (B) Top, time-course of averaged traces and boxplots reporting respectively normalized photons (R = 56 trials, F_3850_= 50.88, ***p<0.0001; AL=23 trials, F_1540_= 3.642, *p=0.012; RM One way ANOVA) and area under the curve (R vs AL, 11.70 ±5.95 vs -12.96 ± 6.85; t_77_=2.40, *p=0.019, Unpaired t-test) for single trials aligned to the behavioral onset. Bottom: same as top but aligned to the offset (R: F_3850_= 65.71, ***p<0.0001; AL: F_1540_= 6.79,***p<0.0001; RM One way ANOVA; AUC analysis: R vs AL, - 42.07 ± 4.01 vs 26.16 ± 5.71; t_77_=9.401, ***p<0.0001, Unpaired t-test). Data are presented with boxplots (median and 10-90 quartile) or mean ± S.E.M.

### Heterogeneity of habenular neuronal activity emerges during defensive behaviors

Analysis of neuronal function with fiber photometry (Figure 2A, B) lacks cellular-level resolution, and provides only aggregated activity from large neuronal populations (Resendez et al., 2016). Such limitation can be circumvented through the use of gradient-refractive-index (GRIN) lenses, which enable visualization of deep-brain neuronal activity with single-cell resolution. We next examined how individual LHb neurons represent threats via their activity patterns. We used a miniature fluorescence microscope to track the relative changes in Ca^2+^ fluorescence in LHb neurons in freely moving mice during threat-driven behaviors (Figure 3A; 62 ± 14.6 neurons per animal; n_mice_ = 4). LHb neurons exhibited diverse activity patterns, with sharp elevations in Ca^2+^ fluorescence during runaway. The response was in the opposite direction during action-locking trials (Figure 3B and Figure 3–figure supplement 1A). The average Ca^2+^ signal across all neurons recorded from a single animal recapitulated the response profiles observed with photometric analysis, supporting the validity of these experimental approaches (Figure 3B, Figure 3–figure supplement 1A and Figure 2B). Thus, single cell analysis of Ca^2+^ signal indicates that opposite neuronal responses in the LHb reflect independent threat-driven behavioral strategies.

**Figure 3.**
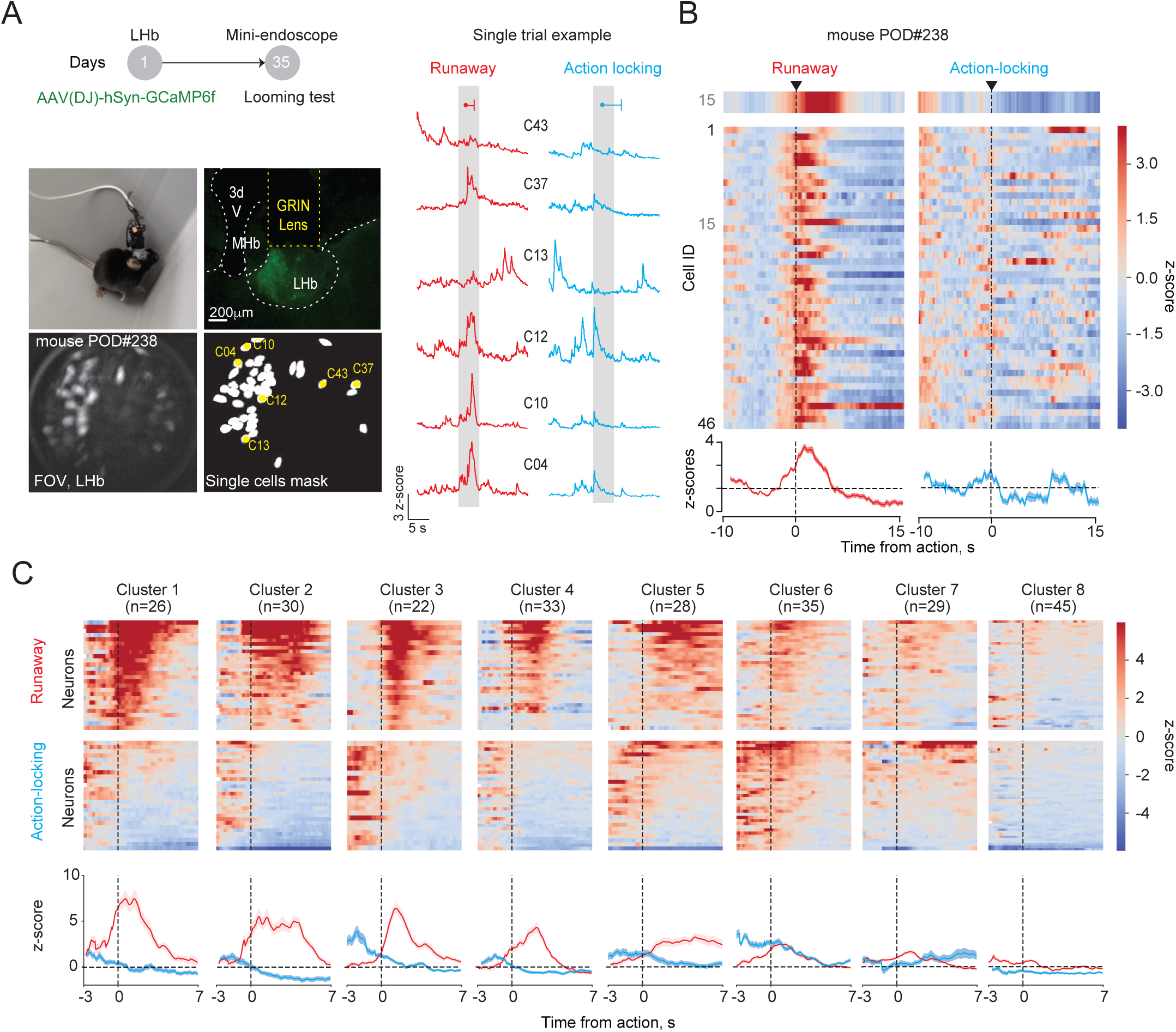
Distinct LHb neuronal ensembles during defensive behaviors. (A) Top: schematic of the experiment. Bottom, pictures showing mouse with miniscope attached, GRIN lens placement, GCaMP6f expression, field of view with identified cells (max intensity projections), map of active LHb neurons and respective sample traces (right). (B) Mean Ca^2+^ responses (z-score) across runaway (left) and action locking (right) trials for 46 LHb neurons imaged within a single mouse, aligned to the onset of the behavioral reaction. Highlighted on the top, the average response of a single cell (Cell ID: 15). Bottom, averaged time-course of all cells for runaway and action locking strategies. (C) Cluster identification by unsupervised classification during runaway (top) and action-locking (center) including all neurons recorded. Bottom, average trace across all neurons within cluster. Plots are aligned to the action onset.

Individual LHb cells displayed variable profiles of runaway-excited/action-locking inhibited responses (Figure 3–figure supplement 1A, B). Furthermore, the activity of single neurons during a given defensive strategy across trials was also variable (Figure 3–figure supplement 1C). Altogether, this argues in favor of functional heterogeneity across LHb neuronal responses after threat. We thereby used an unsupervised clustering algorithm to group the trial-averaged time-locked response of each cell after runaway and action-locking onset (n = 248 from n = 4 mice; Figure 3C and Figure S4A). This analysis revealed eight clusters of neurons based on their responses surrounding the behavioral onset (Figure 3C, Figure 3–figure supplement 2A, B). Clusters were represented in each animal, supporting the strength of independent neuronal representations (Figure 3–figure supplement 2C). The responses of clusters 1 to 5, qualitatively recapitulated fiber photometry Ca^2+^ dynamics time-locked to runaway and action-locking onset (Figure 3C). Cluster 7 and 8, instead, were weakly modulated during looming-triggered defensive responses. Interestingly, Clusters 3 and 6 stood out as their pre-action Ca^2+^ dynamics discriminated the upcoming behavior (Figure 3C). Altogether, this supports the existence of distinct clusters of individual neurons participating throughout threat-driven behavioral responses.

### Decoding the contribution of habenular clusters to threat-driven behaviors

The existence of clusters with neuronal activity that distinguishes the defensive behaviors *prior* to the onset of the action (especially 3 and 6) raised the intriguing possibility that LHb neurons may predict the upcoming selection of runaway or action-locking. To test this idea, we examined the neuronal coding of LHb ensembles by testing whether the defensive strategy on a given trial was identifiable from individual neuron activity patterns (Figure 4A). We defined three time epochs as “prediction of action” (-3 to 0 s from action), “immediate action” (0 to 3 s from action), and “delayed action” (3 to 6 s from action) (Figure 4B). Using leave-one-out cross-validation of a Naïve Bayes classifier (Namboodiri et al., 2019), we calculated the decoding accuracy per neuron above the chance decoding obtained when shuffling trial identity. We then averaged these accuracies across all recorded neurons (Figure 4C) or across all neurons within a cluster (Figure 4D). The null hypothesis was that the average decoding accuracy (above chance) per timeframe and (sub)population is zero. We found that the average decoding accuracy across all recorded neurons was significant for each time epoch (Figure 4C). Interestingly, decoding accuracies showed cluster-specific patterns. Most notably, we found that clusters 3 and 6 showed significant decoding (after correcting for multiple comparison) during the “prediction of action” epoch, whereas other clusters (also including cluster 3) showed significant decoding after the action (Figure 4D). Matching the cluster identity with the topographical neuronal localization during the recordings, revealed that the clusters related to prediction, (clusters 3 and 6), were located caudally with respect to the rest of the clusters (Figure 3–figure supplement 2D).

**Figure 4.**
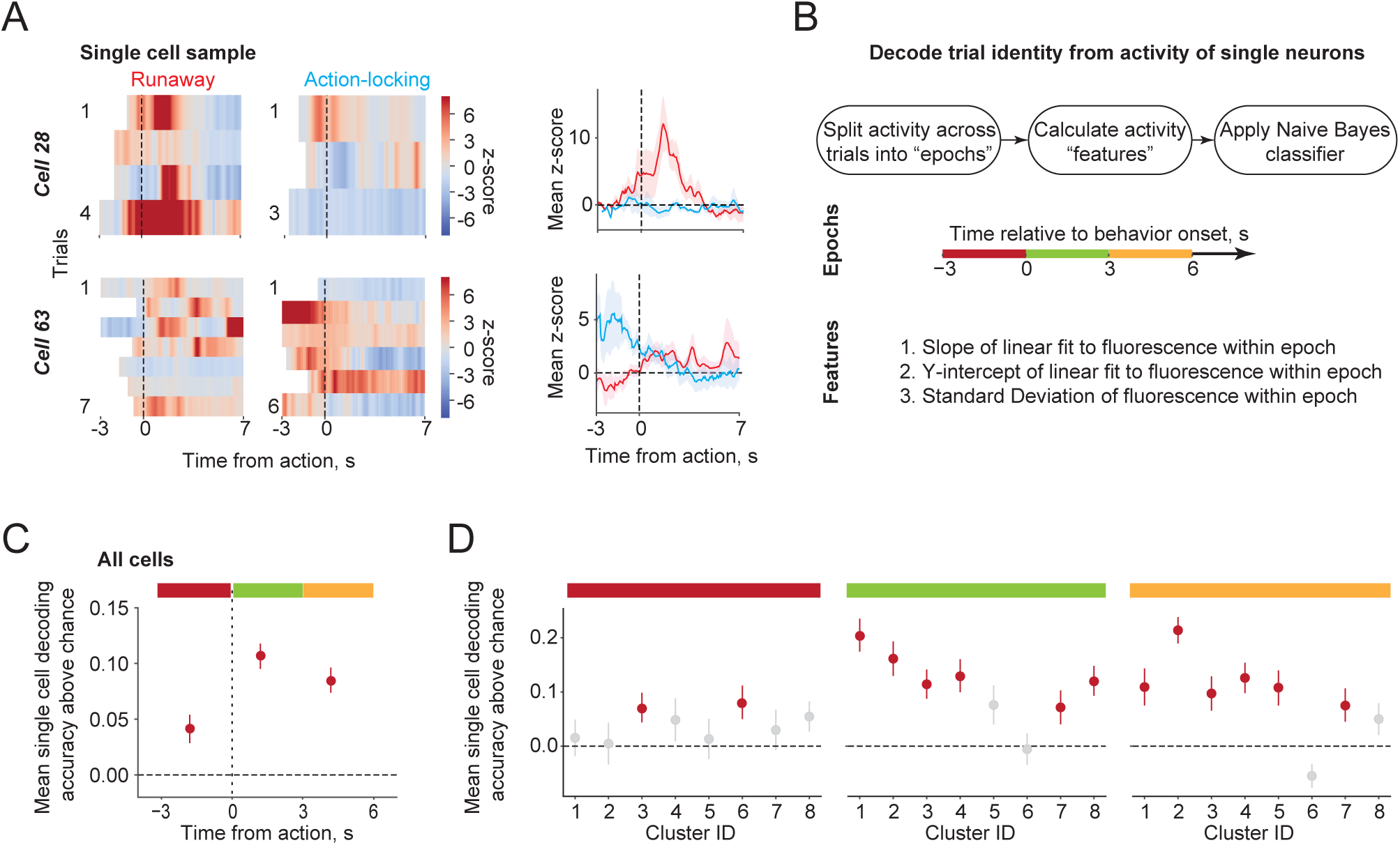
Identified LHb neuronal clusters code for behavioral preparation and execution. (A) Single cell activity across trials during runaway and action locking reported as heat plots (left) and mean z-score (right). Note, trials are time-locked with the behavior and presented different onset due to trial by trial variability in reaction time (blank spaces in the heat plots) (B) Workflow for decoding analysis of single neurons activity. The decoder was run in three different time epochs (-3 to 0 s, burgundy bar; 0 to 3 s, green bar; 3 to 6 s yellow bar) relatively to the behavioral onset. (C) Single cell decoding accuracy above chance averaged acrossall recorded neurons. Red dots highlights significance above chance. Error bars reflect standard error of the mean. t_247_ = 3.23 for -3 to 0 s, t_247_ = 9.37 for 0 to 3 s, t_247_ = 7.54 for 3 to 6 s; p values for the three epochs = 2.67×10^-3^, 1.56×10^-18^ and 6.84×10^-13^ after Benjamini-Hochberg multiple comparisons correction across all epochs. (D) Decoding results split by the clusters. Red dots highlights significance above chance. t_247_ = (0.44, 0.12, 2.54, 1.20, 0.35, 2.58, 0.78, 1.94) for the 8 clusters for -3 to 0 s, t_247_ = (6.62, 5.03, 4.16, 4.20, 2.07, -0.19, 2.25, 4.33) for the 8 clusters for 0 to 3 s, t_247_ = (3.13, 8.66, 2.99, 4.43, 3.27, -2.44, 2.42, 1.67) for the 8 clusters for 3 to 6 s; p values for the three epochs per cluster = (7.55×10^-1^, 9.07×10^-1^, 2.99×10^-2^, 2.96×10^-1^, 7.97×10^-1^, 2.63×10^-2^, 5.28×10^-1^, 7.91×10^-2^) for -3 to 0 s, (2.87×10^-7^, 4.07×10^-5^, 5.35×10^-4^, 3.39×10^-4^, 6.54×10^-2^, 4.43×10^-1^, 4.54×10^-2^, 1.89×10^-4^) for 0 to 3 s, and (7.78×10^-3^, 1.18×10^-10^, 1.12×10^-2^, 1.89×10^-4^, 5.67×10^-3^, 9.84×10^-1^, 3.22×10^-2^, 1.31×10^-1^) for 3 to 6 s after Benjamini-Hochberg multiple comparisons correction across all clusters and epochs.

Overall, these results demonstrate that distinct neuronal subpopulations within the LHb either predict or reflect defensive behavioral selection in response to a threat.

## Discussion

### Dissecting specific contribution of LHb activity for aversion

The past decade witnessed exponentially growing interest in the essential role the LHb has in regulating negatively motivated behaviors. It is of a general consensus within the field that LHb neurons are a homogenous population of glutamatergic cells mostly controlling the function of neuromodulatory systems (Meye et al., 2013). It is also largely accepted that LHb neurons are uniformly excited by aversive external stimuli (Lecca et al., 2017; Matsumoto and Hikosaka, 2007; Wang et al., 2017). Here, we challenge this vision of homogeneity showing that in response to an identical aversive stimulus (the looming), LHb cells dynamics follow opposite logic in a behavior-dependent manner: an escape reaction (runaway) recruits mainly an activation of LHb cells. In contrast, action-locking responses occur along with a decrease in calcium activity, potentially reflecting neuronal inhibition (Namboodiri et al., 2019; Shabel et al., 2019; Wang et al., 2017). Accordingly, aversive foot-shock inhibited neuronal activity of a small and territorially distinct subset of LHb cells (Congiu et al., 2019). Based on this, future work should avoid generalizing that LHb contribution to aversion encoding solely relates to its excitation. Notably, the opposite responses emerging after the looming can occur within the same neuron. It is therefore plausible that a given external stimulus drives dissimilar responses in single cells. The substrate (i.e. connectivity or gene) enabling such neuronal population to encode both behavioral aspects remains however an open question.

### Functional heterogeneity in LHb for threat-driven behaviors

On the basis of recordings and analysis of around 250 LHb cells while animals experience a threat, here we show how ensembles of neurons represent threat-driven behavioral defensive strategies. An unsupervised clustering reveals that independent sets of active neurons form during the expression of threat-mediated behavioral responses (Gründemann et al., 2019; Namboodiri et al., 2019). Such discrete neuronal clusters are stable and define timeframes of threat detection and behavioral action (Gründemann et al., 2019). It remains unclear however which neurobiological substrate defines LHb clusters. Within the amygdala and the cortex, genetically distinct neuronal subtypes contribute to different phases of adaptive behaviors (Abs et al., 2018; Douglass et al., 2017; Krabbe et al., 2019). Recent studies identified molecular-level neuronal diversity within the LHb (Wallace et al., 2019; Hashikawa et al., 2019). Exploiting this genetic knowledge may provide an entry point to specifically probe the functional and behavioral relevance of individual LHb neuronal clusters identified in this study. Alternative to a genetic basis, clusters may emerge according to topographical organization, input-specific connectivity or discrete projection targets (Cerniauskas et al., 2019; Lecca et al., 2017; Meye et al., 2016; Shabel et al., 2012; Valentinova et al., 2019). Our analysis indicates that some LHb neuronal clusters are topographically distinct. This heightens the need of future studies to address this unresolved questions. Notably, the multilevel heterogeneity (functional, anatomical, molecular) emerging lately replaces the initial uniform connotation attributed to the LHb. Further studies will need to determine the relationship across these multiple levels of heterogeneity and establish their behavioral relevance.

### Complex neuronal networks for defensive behaviors

The initial observation that limiting excitation onto LHb impairs escape behaviors implicated this structure in the encoding of innate escape (Lecca et al., 2017). An original aspect of the present work lies on the demonstration that LHb activity changes when animals escape or action-lock after looming presentation. In contrast, recent studies support the contribution of several midbrain nuclei mostly for threat-driven escape (Evans et al., 2018; Seo et al., 2019). Indeed, Ca^2+^ imaging and brain circuit manipulation approaches demonstrate that glutamatergic neurons of the dorsal periaqueductal grey encode decision making and escape (Evans et al., 2018). In addition, a visual pathway engaging superior colliculus and amygdala also contributes to defensive strategies (Shang et al., 2018). Finally, GABAergic neurons in the ventral tegmental area (VTA) projecting to the central amygdala (CeA) seem to be similarly instrumental for threat-driven escape responses (Zhou et al., 2019). Intriguingly, LHb axons innervate these VTA-GABA cells projecting to CeA. Future studies should test how diverse defensive strategies engage wide interconnected networks activity to ultimately build an integrated framework for threat-driven behavioral responses. Defensive strategies are a combination of behavioral sets relying on unique features including trajectories, or stereotyped movements (Evans et al., 2019). The use of deep neural network analysis tracking facets of animal behaviors (Nath et al., 2019; Wiltschko et al., 2015) may pave the way to differentiate precise aspects of defensive behaviors. This will allow a refined alignment with the neuronal dynamics in defined neuronal circuits (Klaus et al., 2017).

The relationship between LHb function and optimal selection of defensive strategies remains correlative after the analysis of the photometric signal. Yet, the unsupervised clustering and decoding analysis support: *i.* that LHb activity codes for distinct behavioral strategies, *ii.* that the dynamics of discrete LHb neuronal clusters reflect precise time epochs of defensive behaviors and *iii*. that these clusters can predict upcoming selection of the action or represent an action itself (Grewe et al., 2017; Namboodiri et al., 2019). Opto or chemogenetic interrogation of LHb neuronal population offers a mean to probe causality between neuronal activity and behaviors (Saunders et al., 2015). However, this intervention is challenging in the present context, as it is limited by the lack of population-specific viral targeting within LHb (i.e. lack of genetic tools for LHb diversity). The manipulation of LHb function in a non cell-specific fashion remains a poor approach to test for causality. This would not fulfill the requirement of precise neuronal cluster targeting, a feature highlighted in the functional and topographical analysis provided in this work. The latest insights of genetic profiling may soon provide the tools to assess these outstanding questions.

In summary, our results identify the evolution of individual neuronal responses in a deep structure like the LHb during threat-driven behavioral strategies, an objective so far proven challenging due to technical difficulties. We demonstrated that LHb neuronal clusters participate to the optimal selection of defensive strategies. Future studies can provide a link between this functional heterogeneity with genetic and anatomical aspects to establish a comprehensive knowledge of LHb contribution to threat encoding. Altogether, these findings advance our understanding of the neuronal basis of ethologically-relevant innate behaviors.

## Contributions

S.L. and M.M. conceptualized the project. S.L. performed and analyzed behaviors and in vivo calcium imaging. L.R. provided support for behavioral analysis and experiments. N.G. performed independent calcium imaging analysis. G.P. provided analytical support for the photometric detection.

V.M.K.N. and G.D.S. provided support, and analysis for calcium imaging analysis and help in editing the manuscript. M.M. and S.L. wrote the manuscript with the help of all authors.

## Acknowledgements

We thank all the members of the Mameli laboratory for comments on the manuscript. We thank C. Lüscher, R. Van Zessen, A. Adamantidis, L. Oesch, J. Zapata, and K. Tan for technical assistance. This work was supported by the ERC StG SalienSy 335333, the Swiss National Funds 31003A and Vaud Canton to M.M., the NARSAD Young Investigator to S.L. and V.M.K.N, and K99MH118422 from US National Institute of Mental Health to V.M.K.N.

## Supplementary figures legends

**Figure 2–figure supplement 1.**
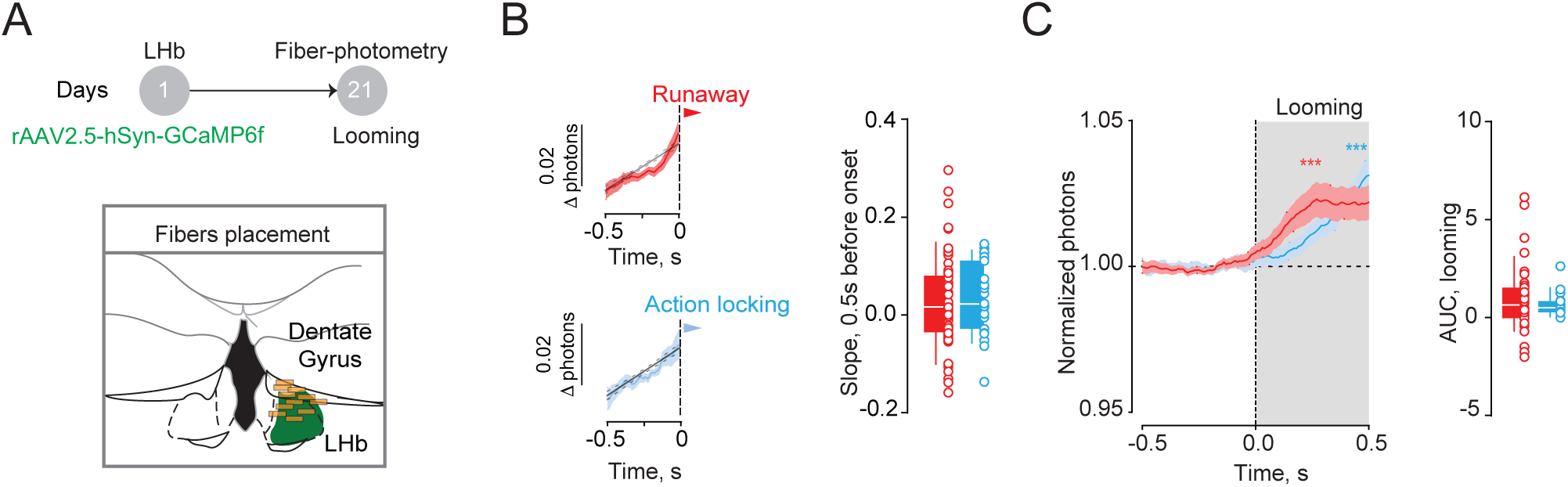
Looming-locked shift in LHb Ca^2+^ signal occurs prior runaway and action-locking. (A) Schematic of fiber placement in the LHb (brown rectangles represent fiber tip placement) (B) Representative averaged traces and boxplots for Runaway (R, 56 trials) and Action-locking (AL, 23 trials) reporting the slope 0.5 s before the behavioral onset (R vs AL, 0.030 ± 0.012 vs 0.032 ± 0.015, t_77_=0.127, p=0.91 Unpaired t-test) (C) Normalized photon traces and area under curve showing the LHb activity time-locked with the looming onset for Runaway (51 trials, F_480_= 4.10; ***p<0.0001, RM One way ANOVA) and Action-locking trials (18 trials, F_153_= 2.45; **p=0.002, RM One way ANOVA). Boxplots reported the AUC for the same set of data (R vs AL, 0.87 ± 0.22 vs 0.63 ± 0.15; t_67_= 0.612, p=0.54 Unpaired t-test). Note that for this analysis trials displaying a behavioral onset < 0.5 sec were discarded to avoid behavior-dependent signal contamination. Data are presented with boxplots (median and 10-90 quartile) or mean ± S.E.M.

**Figure 2–figure supplement 2.**
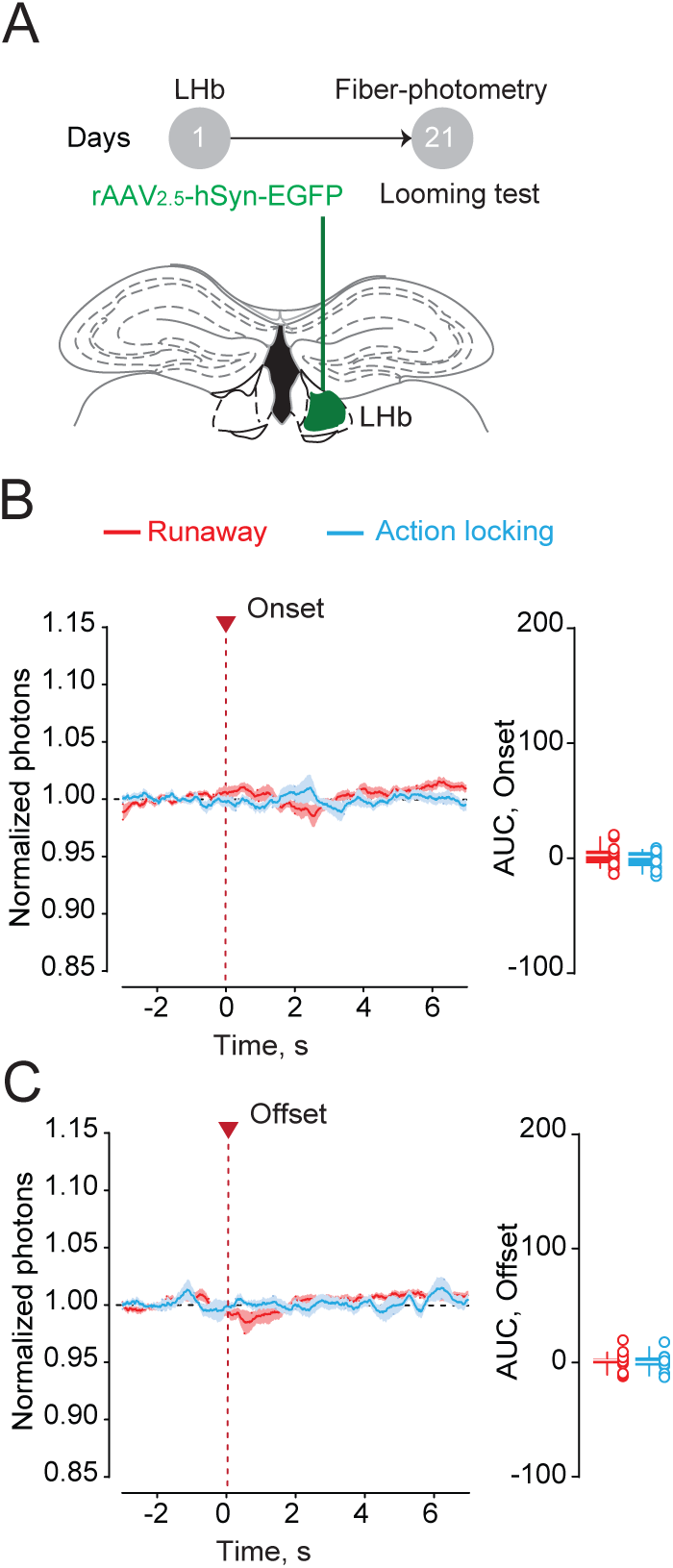
Lack of fluorescent transients in absence of GCamp6f expression. (A) Schematic of the experiment and representative brain coronal section showing eGFP injections in the LHb (n_mice_= 5). (B) Left: Normalized photons time-course graph showing averaged traces of Runaway (R, n_trials_= 21, F_1400_=2.46, p=0.065, RM One way ANOVA) and Action-locking (AL, n_trials_= 12, F_770_=0.946, p=0.401 RM One way ANOVA) trials time-locked with the behavioral onset. Right: area under curve (AUC) for the same data set (R vs AL, 2.662 ± 1.96 vs -0.6096 ± 2.18; t_31_=1.063, p=0.295, Unpaired t-test). (C) Same as (B) but trials are locked with the offset of the behavior (R: F_1400_=0.83, p=0.394; AL: F_770_=0.906, p=0.452; RM One way ANOVA. AUC analysis: (R vs AL, 1.051±1.58 vs 0.3684±2.24; t_31_=0.253, p=0.801 Unpaired t-test). Data are presented with boxplots (median and 10-90 quartile) or mean ± S.E.M.

**Figure 2–figure supplement 3.**
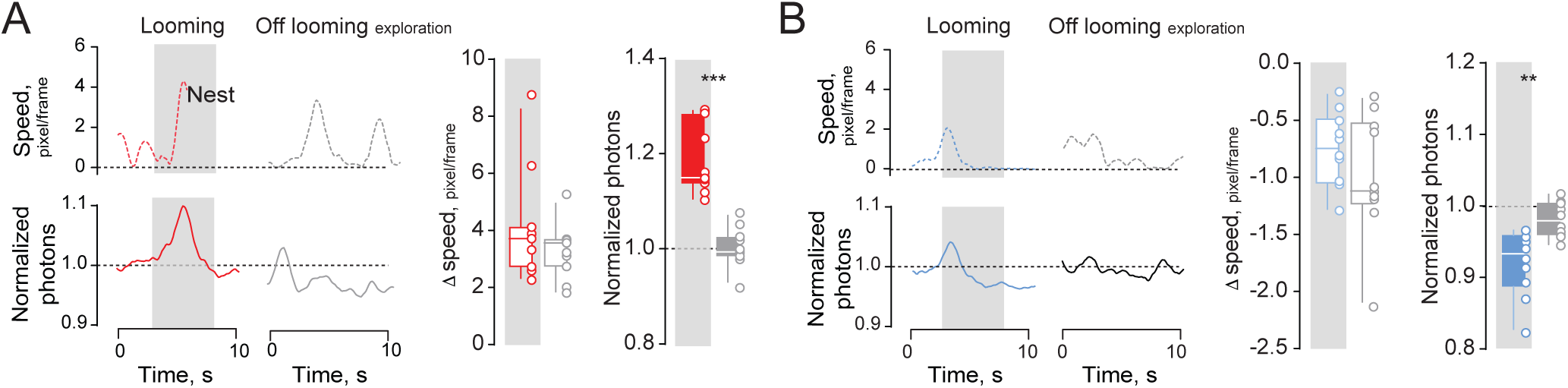
LHb neuronal Ca^2+^ transients are independent of locomotion. (A) Representative traces and boxplots reporting increase in speed (Looming on vs Looming off; 11 vs 11; 4.1 ± 0.56 vs 3.3 ± 0.28 pixel/frame; t_20_=1.28, p=0.21 Unpaired t-test) and the relative LHb activity (Looming on vs Looming off; 11 vs 11, 1.19±0.02 vs 1.00 ± 0.012 normalized photon; t_20_ = 7.35, ***p<0.0001 Unpaired t-test) in presence or absence of the looming stimulus. (B) Same as (A) but for decrease in speed (Looming on vs Looming off; 10 vs 10; -0.7558 ± 0.10 vs -0.9987 ± 0.176 pixel/frame; t_18_=1.190, p=0.249 Unpaired t-test) and relative LHb photon change (Looming on vs Looming off; 10 vs 10, 0.9204±0.014 vs 0.9807±0.007 normalized photons; t_18_=3.61, **p=0.002 Unpaired t-test). For this comparison, we selected the first runaway and action locking response for each mouse. Then we looked for a single episode outside looming presentation with a comparable change in speed for the same mouse. Note that one mouse did not display any action locking response throughout the recording session. Data are presented with boxplots (median and 10-90 quartile) or mean ± S.E.M.

**Figure 3–figure supplement 1.**
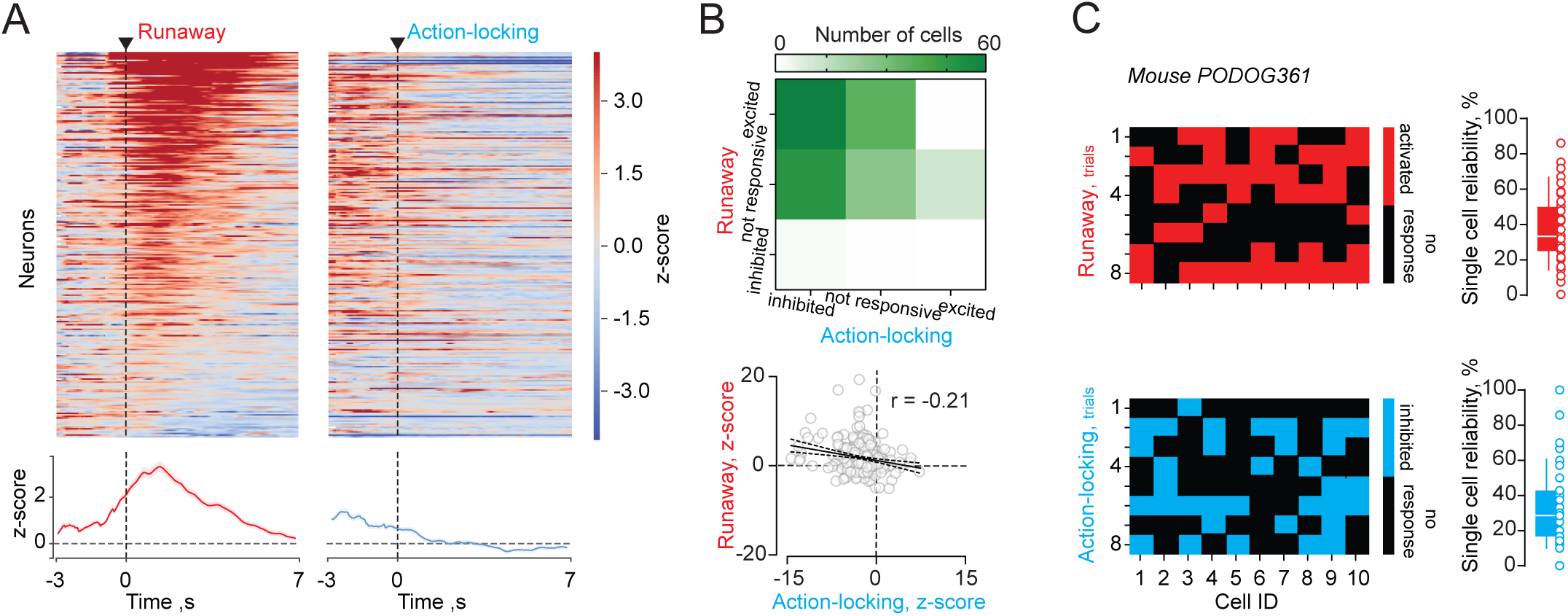
Opposite threat-driven responses occur in the same LHb neuronal ensemble. (A) Mean Ca^2+^ responses for runaway (left) and action-locking (right) trials time-locked with the behavioral onset, including all cells recorded in 4 mice (n=248). Cells are sorted for response magnitude in runaway trials. On the bottom, runaway- and action-locking-locked averaged signals. Data are reported as z-score. (B) On the top, heat-map showing the cell distribution in the different categories according to their response to runaway and action-locking (Runaway/Action-locking: excited/inhibited=73, excited/non responsive=47, excited/excited=4, non responsive/inhibited=62, non responsive/non responsive=35, non responsive/excited=16, inhibited/inhibited=6, inhibited/non responsive=4, inhibited/excited=3). On the bottom, correlation analysis of single cell average Ca^2+^ responses (z-score) to runaway vs action locking displaying variability (Runaway vs Action-locking; n_cells_=248, r=-0.208; R^2^ =0.043; ***p=<0.0001, Pearson correlation coefficient). (C) Top: Raster plots showing active (red squares) and non-active cells (black squares), imaged over different runaway trials in a single mouse. On the right, the boxplot reports single cell reliability (%) for runaway responses (n_cells_=248, Runaway, 38.01 ± 1.3 %). Bottom, same mouse as top. Raster plots showing cells inhibited (red squares) or not (black squares), imaged over different action-locking trials. On the right, the boxplot show reliability for single cells in percentage for action-locking reposes (n_cells_= 248, Action-locking, 32.71± 1.37%).

**Figure 3–figure supplement 2.**
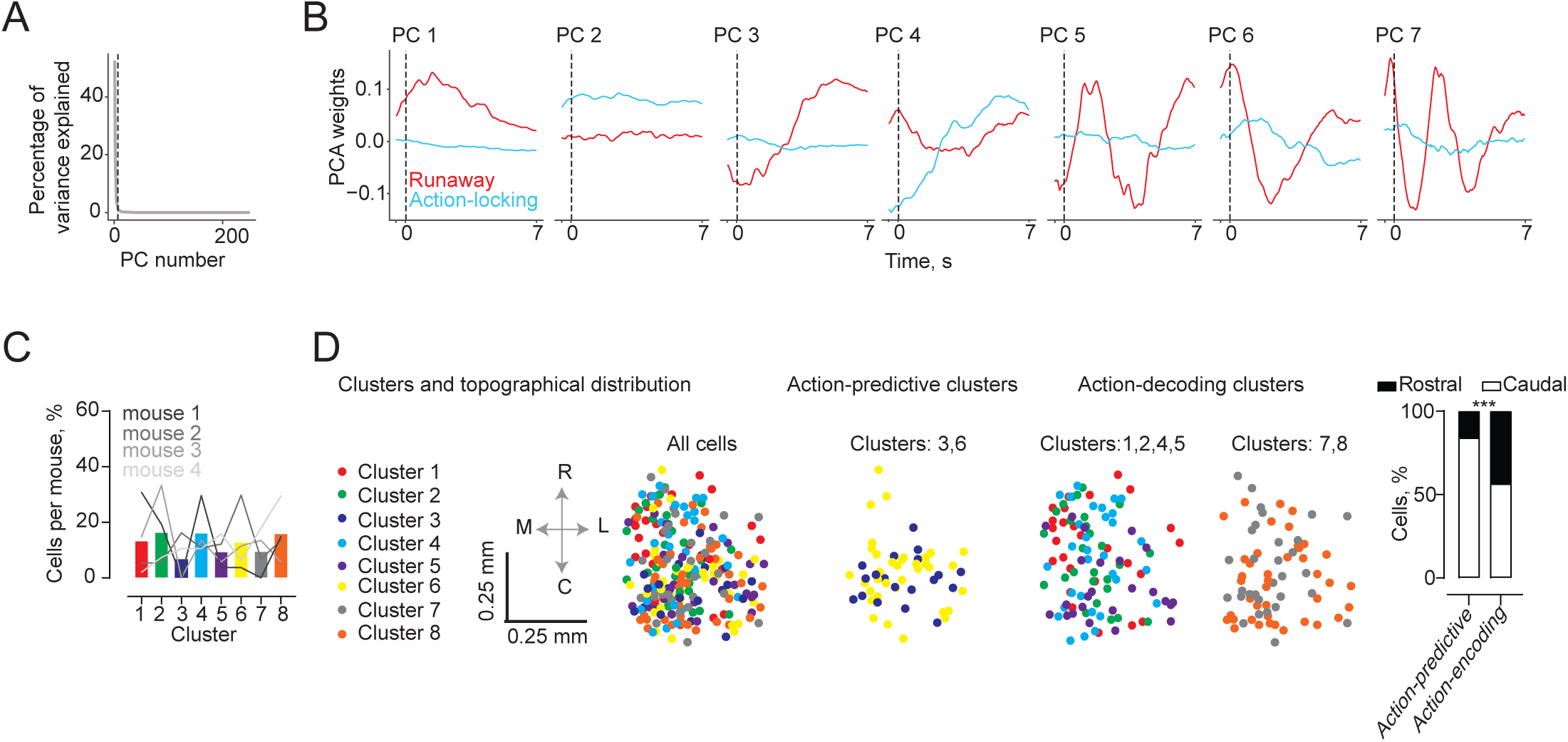
Cluster detection and topography during runaway and action-locking. (A) Plot of the percentage variance explained per principal component, showing the number of principal components retained (dashed line). (B) Individual retained principal components, showing response vectors to both runaway and action-locking trials. (C) The graph reports the percentage of cells in each cluster (Cluster 1 to 8, number of cells per cluster: 26, 30, 22, 33, 28, 35, 29, 45) and single-mouse contribution per cluster (Cluster 1 to 8, number of cells per cluster. Mouse 1: 16, 10, 0, 7, 3, 6, 7, 3. Mouse 2: 4, 4, 12, 8, 9, 22, 5, 10. Mouse 3: 4, 9, 0, 8, 1, 1, 0, 4. Mouse 4: 2, 7, 10, 10, 15, 6, 17, 28). (D) Topographical distribution of the clusters in LHb (action-predictive vs action-decoding clusters; rostral vs caudal cell distribution; Action-predictive: 57 cells, 9 rostral vs 48 caudal. Action-decoding: 191 cells, 83 rostral vs 108 caudal. X^2^1= 14.4; z=3.79; ***p=0.0001, Chi-Square test).

## Material and methods

### Experimental subjects

The experiments were performed on C57Bl/6J mice wild-type males of 10-18 weeks. Mice were housed at groups of five per cage with water and food ad libitum on a 12:12 h light cycle (lights on at 7 a.m.). All procedures aimed to fulfill the 3R criterion and were approved by the Veterinary Offices of Vaud (Switzerland; License VD3171).

### Behavioral paradigm

Mice were tested for behavior in a looming visual stimulus test, as described elsewhere (Yilmaz and Meister, 2013). Animals were placed in an open-top plexiglas box (58cm L× 38cm W× 32cm H). A triangular shaped nest (20 × 12 cm) was placed in one corner. Recordings were performed under illumination provided by the projector screen (52 cm × 30 cm; Dell) and an infrared light-emitting diode (LED) illuminator (Pinnacle Technology), both placed above the arena. Experiments were recorded at 60 frames per second with a near-IR GigE camera (acA1300-60gmNIR, Basler) positioned in one side of the arena. Video recording, was controlled with Ethovision and synchronized with the photometric and endoscopic recordings using hardware-time signals controlled with a I/O box (Noldus). All the mice tested underwent a period of habituation to the fiber/camera spanning from 15-20 min session every day for 3 consecutive days. For the experiment, after 5-10 min of acclimatization, a looming stimulus (always delivered at 50% contrast), was randomly presented from the screen in the center of the arena while the mouse was actively exploring (independently by its position in the arena). The stimulus of 0.5 s duration was repeated 5 times with an inter-stimulus interval of 0.5 s. Each mouse received from 7 to 20 trials with a minimum inter-trial interval of about 5 minutes. The video analysis of the behavior was performed off-line.

#### Automated detection of mouse shape and position

A fully convolutional neural network was used to extract the shape of the mouse across the arena. Each video (1920 x 1088 @ 60 fps) was converted to a sequence of images (8-bit, 256 x 144 pixel). The training dataset was composed of 112 images and it was used to trace a set of 112 masks (8-bit, 256 x 144 pixel binary images) delineating the contour of the mouse body and to output the files storing the coordinates of the center of mass of each individual mask. Each image in the training dataset was passed through three convolutional layers (channels: 16, 32, 64; kernels: 3, 5, 3, stride: 1, ReLU units), two max-pooling operations (kernel size: 2), and three transposed convolutional layers. The frames were processed in batches of 64 images for 171 epochs). The network was built with the open source library PyTorch 1.2 (https://pytorch.org/) and trained to minimize (Adam optimizer, learning rate: 0.003) the Mean-Squared Error loss function. Accuracy was measured as the Euclidean distance between the centroid of the mask of the training set and the centroid of the score map predicted by the network. An arbitrary cutoff was used to define the boundaries of the estimated mouse shape on the score map. The mean accuracy on the test set was 1.65 px (+/- 1.51 px, standard deviation), with 96.4% of the frames showing a distance between centroids (i.e. label Vs predicted) less than 7 px. The output coordinates of the center of mass were then used to compute the speed (pixels/seconds) and the location of the mouse inside the arena. The onset of runaway was measured as the peak of the first derivative of the mouse speed tracking curve. The runaway offset was coinciding with the mouse entrance in the nest. The score map was used to estimate the size of the mouse (e.g. total number of pixel above the arbitrary threshold) across the arena and used for further calculations to score action-locking behavior.

#### Automated classification of action-locking behavior

An observer blind to the experimental condition of the animals manually scored the action locking behavior, defined as a sudden blockade of all -except respiratory-movements. In contrast to freezing, action locking was not associated with a particular body posture (i.e. crouching). The sudden immobility had to last at least two seconds in order to score the animal as actively producing an action-locking behavior. Data obtained from the manually labeled frames were then merged with the data (speed and size) obtained from the automatic detection of the mouse position to train a random forest classifier to predict in each frame whether the animal was in action-locking. Both speed and size were convolved with a Max function (window = 60 frames) and a total of four features were used: speed (v), size (s), es, and ev. A 5-fold cross-validation yielded an overall accuracy of 98%. The accuracy achieved on the test set was 97.5% with a false positive rate of 2.6%.

### Surgical procedures

#### Viral injections

All mice were anaesthetized with ketamine (150 mg/kg)/xylazine (10 mg/kg) (Sigma-Aldrich, France). We unilaterally injected in the LHb (-1.4 mm AP, 0.45 ML, 3.1 mm DV) rAAV2.1-hSyn-GCaMP6f-eGFP or rAAV/DJ-hSyn--GCaMP6f-eGFP or rAAV2.5-hSyn-eGFP (University of North Carolina, US) using a glass pipette on a stereotactic frame (Kopf, France). Volumes ranged between 200 and 300 nl, at a rate of approximately 100-150 nl/min. The injection pipette was withdrawn from the brain 10 minutes after the infusion. Animals were allowed to recover for a minimum of two weeks before fiber or GRIN lenses implantation.

#### Chronic implants

For fiber photometry experiments, a single fiber probe was placed and fixed (C and B Metabond, Parkell) 150 μm above the injection site in isoflurane anesthetized (induction: 4%, maintenance: 1.8-2%) mice. For endoscope experiments, mice were anaesthetized (as described above) and implanted with a GRIN (Graded-Index) lens (6.1mm length, 0.5mm diameter; Inscopix, #100-000588). The lens was targeted to be ∼ 150–200 μm above the injection site using the following coordinates: −1.40 mm posterior to bregma, 0.45 mm lateral from midline, and −2.85 to −2.9 mm ventral to skull surface (lowered at a speed of 1µm/s). To increase stability of the implants the lenses were implanted into the dorsal portion of the region allowing imaging ventral LHb neurons. Two week after lens implantation, mice were again anaesthetized (isoflurane, as above) and a baseplate (Inscopix, #100-000279) was secured above the lens. A baseplate cover (Inscopix, #100-000241) was attached to prevent damage to the microendoscope lens. Out of 23 mice that were injected with GCaMP6f virus, 4 had successful lens implantation/viral expression and were used for this study.

### Fiber photometry recordings

Fiber photometry measurements were carried out by the ChiSquare X2-200 system (ChiSquare Biomaging, Brookline, MA). Briefly, blue light from a 473-nm picosecond-pulsed laser (at 50 MHz; pulse width ∼ 80 ps FWHM) was delivered to the sample through a single mode fiber. Fluorescence emission from the tissue was collected by a multimode fiber with a sample frequency of 100Hz. The single mode and multimode fibers were arranged side by side in a ferrule that is connected to a detachable multimode fiber implant. The emitted photons collected through the multimode fiber pass through a bandpass filter (FF01-550/88, Semrock) to a single-photon detector. Photons were recorded by the time-correlated single photon counting (TCSPC) module (SPC-130EM, Becker and Hickl, GmbH, Berlin, Germany) in the ChiSquare X2-200 system.

### Endoscope recordings

All calcium imaging was recorded at 20 frames per second, 200-ms exposure time, and 10–40% LED power (0.4-0.9mW at the objective, 475nm) using a miniature microscope from Inscopix (nVista). Calcium recording files were down-sampled (spatial binning factor of 4) to reduce processing time and file size, filtered, corrected for rigid brain movement and the ΔF/F0 was calculated using as F0 the average fluorescence for all the video (Inscopix, IDP). Individual component analysis and principle component analysis (ICA/PCA) applications were used to identify individual cells and to extract their respective calcium traces.

In addition, to compare ROI detections and relative traces obtained with the PCA/ICA we also performed constrained non-negative matrix factorization for endoscopic data (CNMF-E) for a subset of data. Briefly, we denoised, deconvolved, and demixed calcium-imaging dynamics (http://www.github.com/zhoupc/cnmf_e). This method allows accurate single neurons fluorescence traces extraction (Zhou et al. 2018). Calcium imaging frames were initially pre-processed in Mosaic (Inscopix) for motion correction. We use a Gaussian kernel width 4 μm, maximum soma diameter 16 μm, minimum local correlation 0.8, minimum peak-to-noise ratio 8 and merging threshold was set to 0.65 for optimal discrimination of temporal and spatial overlap.

### Analysis

Photometric signal as well as ICA/PCA derived traces were smoothed (constant time factor, 0.1 s) and further processed according to the trials using Spike2 software (Cambridge Electronic Design). We obtained an average peri-stimulus time histogram (PSTH) trace aligned to the stimulus or behavioral onset/offset (3 s prior and 7 sec after a given event). For the photometric recordings we calculate the photon change normalizing for the 3 sec prior each trial. For the endoscope recordings we z-scored each trials in reference to their baseline (3 s prior to behavior onset).

We identified functional sub-classes of neurons by comparing the fluorescence Ca2+ signals of individual cells before and after a given event, using 2s time span. For runaway trials we consider a cell excited if the signal 2 s post runaway onset was higher than the baseline plus 2 SD. Vice versa a cell was inhibited if its signal in the 2s post runaway resulted 2 SD lower than their baseline. For action-locking responses we considered 3 epochs (2s each epoch) of analysis post event according with the average duration of this behavior (6s). If the signal in at least one epoch resulted higher or lower than 2 SD of the baseline the cell was considered action-locking excited or inhibited respectively.

For the analysis of the single trials we follow the same logic above-mentioned except that the epochs considered for the action locking were updated each time according with the duration of the response.

### Clustering and decoding

For clustering neurons based on their average responses around action onset for both action-locking and runaway trials, we followed a similar general procedure as in Namboodiri et al. 2019. Briefly, we first calculated the average peri-event time histogram (PETH) for each neuron around each action by averaging all trials. Due to the variability in reaction times from looming stimulus onset until the action, we calculated the PETHs around a time window from -0.5 s to +7s surrounding the action. This ensured that only activity after the looming stimulus onset was included in all trials. The PETH surrounding both action-locking and runaway trials were treated as features of the response of a neuron. This feature space was then reduced in dimensionality using principal components analysis (Fig S5). The number of principal components to keep was decided based on the bend in the scree plot (Namboodiri et al. 2019). A spectral clustering algorithm along with optimal selection of number of clusters using silhouette scores (Namboodiri et al. 2019) was used on the principal component scores to test for presence of clusters. The number of clusters was chosen by maximizing the silhouette score. Once cluster identities were assigned, all PETHs were recalculated using the activity from -3 s to +7 s surrounding the actions. Only activity following looming stimulus onset was included. If the looming stimulus onset was less than 3 s prior to action on a trial, these data were treated as “not a number (nan)” in our analysis pipeline.

We then tested for significant decoding by analyzing whether the activity of a single neuron could be used to decode the chosen behavioral action on a trial. To calculate a decoding accuracy, we trained a Naïve Bayes classifier on all but one trial (leave-one-out cross-validation) and tested the decoding accuracy on the remaining trial for each time epoch (Figure 4B). Within each epoch, three “response features” were used for decoding analysis: slope of the linear fit to fluorescence within the epoch, y-intercept of this fit, and lastly, the standard deviation of fluorescence within the epoch (Figure 4B). Only three features were used to avoid overfitting and maximize generalizability of decoding on test trials. This procedure was repeated with each trial as the test trial, to obtain an overall decoding accuracy above chance accuracy obtained by shuffling trial identity. For the shuffled null, we calculated the mean chance accuracy per neuron as the mean accuracy across ten different shuffles. We applied this procedure to one neuron at a time to obtain a decoding accuracy per neuron, which was then averaged across all neurons recorded, or all neurons within a cluster. The decoding accuracy above chance was simply calculated as the difference in population mean between the true accuracies and the shuffled accuracies. Significance was tested based on a two-sample t-test between the true accuracies and the shuffled accuracies.

### Statistical analysis

Offline analyses were performed using Prism 8 (Graphpad, US). Single data points are always plotted. Sample size was pre-estimated from previously published research and from pilot experiments performed in the laboratory. Each mouse represents an analytical unit, for each experiment we stated the replication factor. Compiled data are expressed as boxplots (median and quartiles) or mean ± S.E.M. Significance was set at p < 0.05 using two-sided unpaired t-test, one or two-way ANOVA. Correlational analysis was performed with Pearson test. Frequency distribution was analyzed with Χ^2^ test. The use of the paired t-test and two way ANOVA for repeated measured were stated in the legend figure text.

